# Rapid Vacuolar Sorting of the Borate Transporter BOR1 Requires the Adaptor Protein Complex AP-4 in Arabidopsis

**DOI:** 10.1101/2023.12.06.570355

**Authors:** Akira Yoshinari, Yutaro Shimizu, Takuya Hosokawa, Akihiko Nakano, Tomohiro Uemura, Junpei Takano

## Abstract

Plants maintain nutrient homeostasis by controlling the activities and abundance of nutrient transporters. In *Arabidopsis thaliana*, the borate (B) transporter BOR1 plays a role in the efficient translocation of B under low-B conditions. BOR1 undergoes polyubiquitination in the presence of sufficient B and is then transported to the vacuole via multivesicular bodies (MVBs) to prevent B accumulation in tissues at a toxic level. A previous study indicated that BOR1 physically interacts with μ subunits of adaptor protein complexes AP-3 and AP-4, both involved in vacuolar sorting pathways. In this study, we investigated the roles of AP-3 and AP-4 subunits in BOR1 trafficking in Arabidopsis. The lack of AP-3 subunits did not affect either vacuolar sorting or polar localization of BOR1-GFP, whereas the absence of AP-4 subunits resulted in a delay in high-B-induced vacuolar sorting without affecting polar localization. Super-resolution microscopy revealed a rapid sorting of BOR1-GFP into AP-4-positive spots in the *trans*-Golgi network (TGN) upon high-B supply. These results indicate that AP-4 is involved in sequestration of ubiquitinated BOR1 into a TGN-specific subdomain “vacuolar-trafficking zone,” and is required for efficient sorting to MVB and vacuole. Our findings elucidate the rapid vacuolar sorting process facilitated by AP-4 in plant nutrient transporters.

**Subject areas:** (7) Membrane and transport

## Introduction

Plants regulate nutrient uptake and translocation through a variety of transport proteins. The abundance of these nutrient transporters is regulated through multiple steps, including RNA transcription, protein translation, and post-Golgi trafficking processes (i.e., secretion, endocytosis, recycling, and vacuolar transport) (Paez Valencia et al., 2016; Gu et al., 2017; Ivanov and Vert, 2021; González Solís et al., 2022). In plants, specific mineral transporters located in the plasma membrane (PM), such as IRON-REGULATED TRANSPORTER 1 (IRT1), PHOSPHATE TRANSPORTER 1 (PHT1), and REQUIRES HIGH BORON 1 (BOR1), undergo ubiquitination and subsequent vacuolar sorting for degradation in response to an increase in their substrate concentration (Takano et al., 2005; Kerkeb et al., 2008; Bayle et al., 2011; Barberon et al., 2011; Kasai et al., 2011; Huang et al., 2013). However, the precise intracellular trafficking routes of the transporters to the vacuole after ubiquitination have not yet been fully elucidated.

The boric acid/borate transporter BOR1 is required for the normal growth of Arabidopsis under low-boron (B) conditions (Takano et al., 2002). In the PM, BOR1 exhibits polar localization towards the stele (inner-lateral side) in various root cells under low-B conditions (Takano et al. 2010; Yoshinari et al. 2016). Alongside the boric acid channel NIP5;1, which exhibits a polar localization in the PM towards the soil side (outer lateral side) in root epidermal and cortical cells, BOR1 transports B to the center of the roots (Yoshinari and Takano, 2017; Jothi and Takano, 2023). Under low-B conditions, the cytoplasmic fraction of BOR1 is preferentially localized in the *trans*-Golgi network (TGN), which functions as an early endosome (EE) in plants, and is endocytosed and recycled between the PM and TGN (Takano et al., 2005; Yoshinari et al., 2018). In contrast, upon high-supply, BOR1 is rapidly ubiquitinated and subsequently transported into the vacuole for protein degradation, via the multivesicular body/late endosome (MVB/LE) (Takano et al., 2005, 2010; Viotti et al. 2010; Kasai et al., 2011; Yoshinari et al., 2021a). The B-induced vacuolar sorting of BOR1 requires predicted sorting signals in its cytosolic large-loop region, such as tyrosine motifs (YxxΦ) and dileucine motif ([D/E]xxxL[L/I]) (Takano et al., 2010; Kasai et al., 2011; Wakuta et al., 2015). These sorting signals are involved in the vesicle transport mediated by adaptor protein (AP) complexes (AP-1 to AP-4). AP complexes consist of two large subunits (α, β, γ, ε), one medium subunit (μ), and one small subunit (σ), and are conserved in eukaryotes, including plants (Hirst et al., 1999). A yeast two-hybrid assay indicated that μ subunits of AP-3 and AP-4 can bind to the tyrosine motifs of BOR1, suggesting potential involvement of AP-3 and AP-4 in the intracellular trafficking of BOR1 (Yoshinari et al., 2019).

However, the role of the AP-3 in vesicle formation remains unclear in plant cells. While it is associated with endomembrane compartments, it is unlikely to be localized in either the TGN or the MVB (Feraru et al., 2010). Instead, it appears to be essential for the trafficking pathway to bypass the conventional MVB-to-vacuole pathway (Feraru et al., 2010; Zwiewka et al., 2011). AP-3 is required for the biogenesis and function of lytic vacuoles rather than for protein sorting to storage vacuoles (Feraru et al., 2010; Zheng et al., 2019). In addition, AP-3 is required for vacuolar transport of the tonoplast-localized proteins PROTEIN S-ACYL TRANSFERASE 10 (PAT10) and SUCROSE TRANSPORTER 4 (SUC4) (Feng et al., 2017; Zheng et al., 2019). In the mutants of AP-3, PAT10 and SUC4 are retained in the Golgi apparatus, suggesting that AP-3 mediates the Golgi-to-vacuole transport of membrane proteins (Feng et al., 2018). The role of AP-4 in vacuolar transport in plant cells has been previously reported (Fuji et al., 2016; Müdsam et al., 2018; Shimizu et al., 2021). AP-4 is localized in the TGN and plays a crucial role in recruiting an EPSIN-like accessory protein, MODIFIED TRANSPORT TO THE VACUOLE 1 (MTV1), which facilitates vesicle biogenesis for vacuolar transport (Heinze et al., 2020). Alongside VESICLE-ASSOCIATED MEMBRANE PROTEIN 727 (VAMP727), a subtype of the soluble N-ethylmaleimide-sensitive factor attachment protein receptor (SNARE), AP-4 localizes to a specific subdomain within the TGN known as “vacuolar-trafficking zone” (Shimizu et al., 2021). The lack of AP-4 subunits results in the mislocalization of vacuolar storage proteins, such as 12S globulin, and tonoplast-localized proteins, such as MOLYBDATE TRANSPORTER 2 (MOT2), NATURAL RESISTANCE-ASSOCIATED MACROPHAGE PROTEIN 3 (NRAMP3), and NRAMP4 (Fuji et al., 2016; Müdsam et al., 2018). Accumulating evidence indicates that both AP-3 and AP-4 are involved in vacuolar transport of membrane proteins. However, it remains uncertain whether AP-3 and AP-4 play a role in the vacuolar sorting of ubiquitinated cargo proteins, including mineral transporters, in plants. In this study, we evaluated the involvement of AP-3 and AP-4 in the B-dependent degradation of BOR1 in *Arabidopsis thaliana* and revealed the function of AP-4 in BOR1 sorting at the TGN subdomain.

## Results

### AP-3 and AP-4 are not required for polar localization of BOR1

In our previous study, AP-3 and AP-4 showed the potential to physically bind to the tyrosine motifs of BOR1, which are essential for polar localization and vacuolar sorting of BOR1 (Takano et al., 2010; Yoshinari et al., 2019). To investigate the functional importance of AP-3 and AP-4 in the polar localization of BOR1, we performed confocal microscopy of BOR1-GFP in the primary root tips of mutants lacking the single subunits of AP-3 and AP-4 (*ap3m-1, ap3b-1, ap4m-3*, and *ap4m-4*) (Niihama et al., 2009; Feraru et al., 2010; Fuji et al., 2016) (Fig. 1). We quantified the polarity index of BOR1-GFP in the wild-type and mutants grown under a low-B condition (0.5 μM) by measuring the GFP fluorescence within the inner and outer half of the PM as previously described (Figs. 1B and 1D) (Yoshinari and Takano, 2020). We noticed that the polarity indexes using the BOR1-GFP construct without the 5’ untranslated region (5’UTR), which is responsible for the B-dependent translational repression of BOR1 (Aibara et al., 2018), is slightly lower than that in 5’UTR for unknown reasons (Fig. 1B; Supplementary Fig. S1). Therefore, statistical analyses for *ap3b-1* and *ap3m-1* were performed separately (Fig. 1B). The polarity indices of BOR1-GFP in all mutants were comparable to those in wild-type plants (Figs. 1B and 1D), indicating that the polar localization of BOR1 requires neither AP-3 nor AP-4 function. The polar localization of BOR1 is dependent on continuous endocytosis and endosomal recycling (Yoshinari et al. 2016, 2019). The lack of effect on the polar localization of BOR1 is consistent with previous reports suggesting that AP-3 and AP-4 play roles in vacuolar transport pathways rather than endocytosis and recycling pathways (Feraru et al., 2010; Heinze et al., 2020; Shimizu et al., 2021; Kleine-Vehn et al., 2011).

**Fig. 1.**
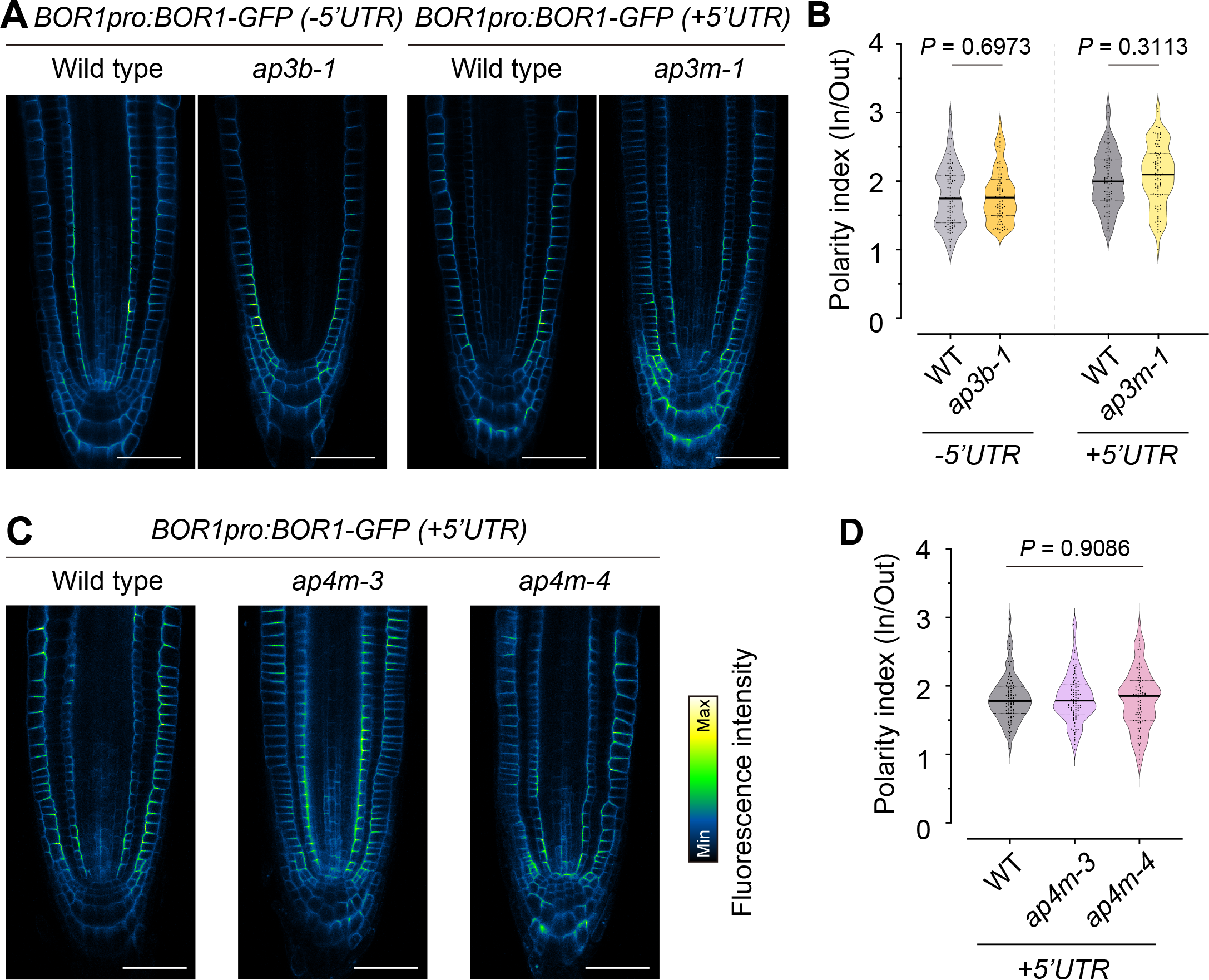
Neither AP-3 nor AP-4 contribute to polar localization of BOR1. Analysis of the polar localization of BOR1-GFP in plants grown under low-B condition (0.5 μM). (A) Confocal images of BOR1-GFP (*proBOR1:BOR1-GFP* ±5’UTR) in wild-type (WT), *ap3b-1*, and *ap3m-1*. (B) Violin plots of polarity index of BOR1-GFP in epidermal cells. *n =* 84 (-5’UTR, WT), 82 (-5’UTR, *ap3b-1)*, 84 (*+*5’UTR, WT), and 84 (*+*5’UTR, *ap3m-1*) cells, respectively, from 7 different plants. *P* values were determined using the Student’s two-tailed *t-*test. (C) Confocal images of BOR1-GFP (*proBOR1:BOR1-GFP* +5’UTR) in WT, *ap4m-4*, and *ap4m-4*. (D) Violin plots of polarity index of BOR1-GFP in epidermal cells. *N =* 84 (WT), 98 (*ap4m-3)*, and 84 (*ap4m-4*) cells from six or seven different plants. *P* value was determined using a one-way ANOVA. Scale bars represent 50 μm.

### Rapid vacuolar sorting of BOR1 requires AP-4 but not AP-3

Subsequently, we assessed the involvement of AP-3 and AP-4 in vacuolar sorting of BOR1. To achieve this, BOR1-GFP was monitored in the *ap3m* and *ap4m* mutants after shifting form a low-(0.5 μM) to high-B condition (100 μM) through confocal microscopy. We measured the fluorescence intensity of BOR1-GFP in the PM of root epidermal cells and compared the rate of BOR1-GFP removal from the PM in the wild-type and mutants. After high-B supply, the PM fraction of BOR1-GFP rapidly decreased in the wild-type, as previously reported (Figs. 2A–2C; Supplementary Fig. S2) (Yoshinari et al., 2021a). BOR1-GFP was more stable in the PM of *ap4m-3, ap4m-4*, and *ap4b-2* mutants than in the wild-type (Figs. 2A–2C). Relative fluorescence intensity of BOR1-GFP in the PM was significantly higher in the *ap4m-3* (0.29±0.02), *ap4m-4* (0.23±0.01), and *ap4b-2* (0.23±0.01) mutants than in the wild-type (0.09±0.01) at 2 h after high-B supply (Fig. 2C). In addition, the rate of BOR1-GFP degradation in *ap3m-1* and *ap3b-1* cells was not significantly affected (Fig. 3), indicating that AP-3 function is dispensable for rapid vacuolar sorting of BOR1. These data indicate that the efficient vacuolar transport of BOR1 requires AP-4, but not AP-3.

**Fig. 2.**
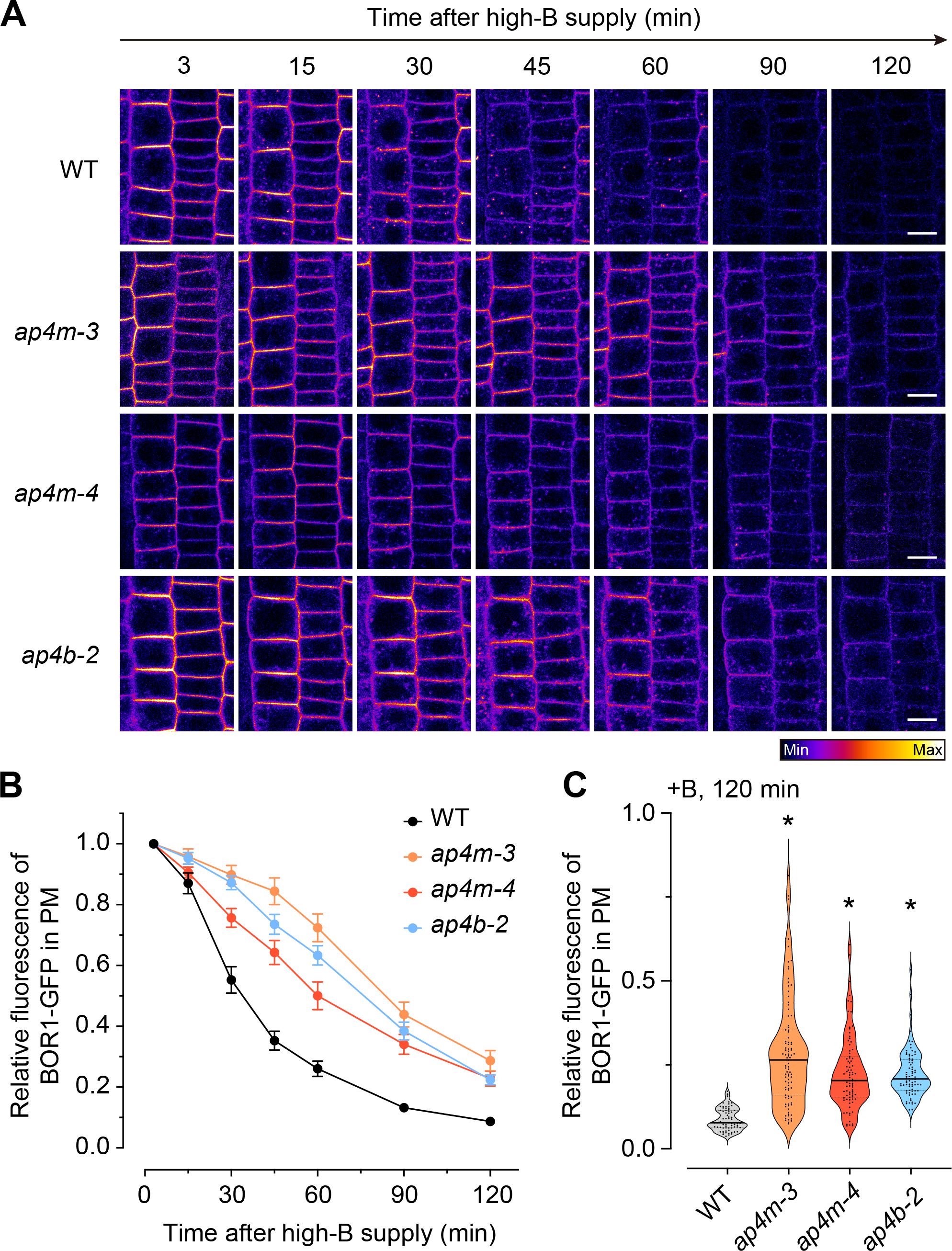
Boron-induced rapid vacuolar sorting of BOR1 requires AP-4 function. Time-lapse imaging of BOR1-GFP in the root epidermis after high-B supply. (A) Confocal images of BOR1-GFP in the wild-type, *ap4m-3, ap4m-4*, and *ap4b-2*. Four-day-old seedlings grown on a medium containing 0.5 μM boric acid were gently put on a glass-bottom dish and underlaid with a soft medium containing 100 μM boric acid. Scale bars represent 10 μm. (B) Fluorescence intensity of BOR1-GFP in the PM. The relative values were calculated by setting the first 3 min to 1. *n =* 71 (WT), 95 (*ap4m-3*), 84 (*ap4m-4*), and 78 (*ap4b-2*) cells from three or four different plants. (C) Violin plots representing individual data points at 120 min (B). Asterisks indicate *P* < 0.0001 as determined by one-way ANOVA with Dunnett’s post-hoc test compared to the wild-type.

**Fig. 3.**
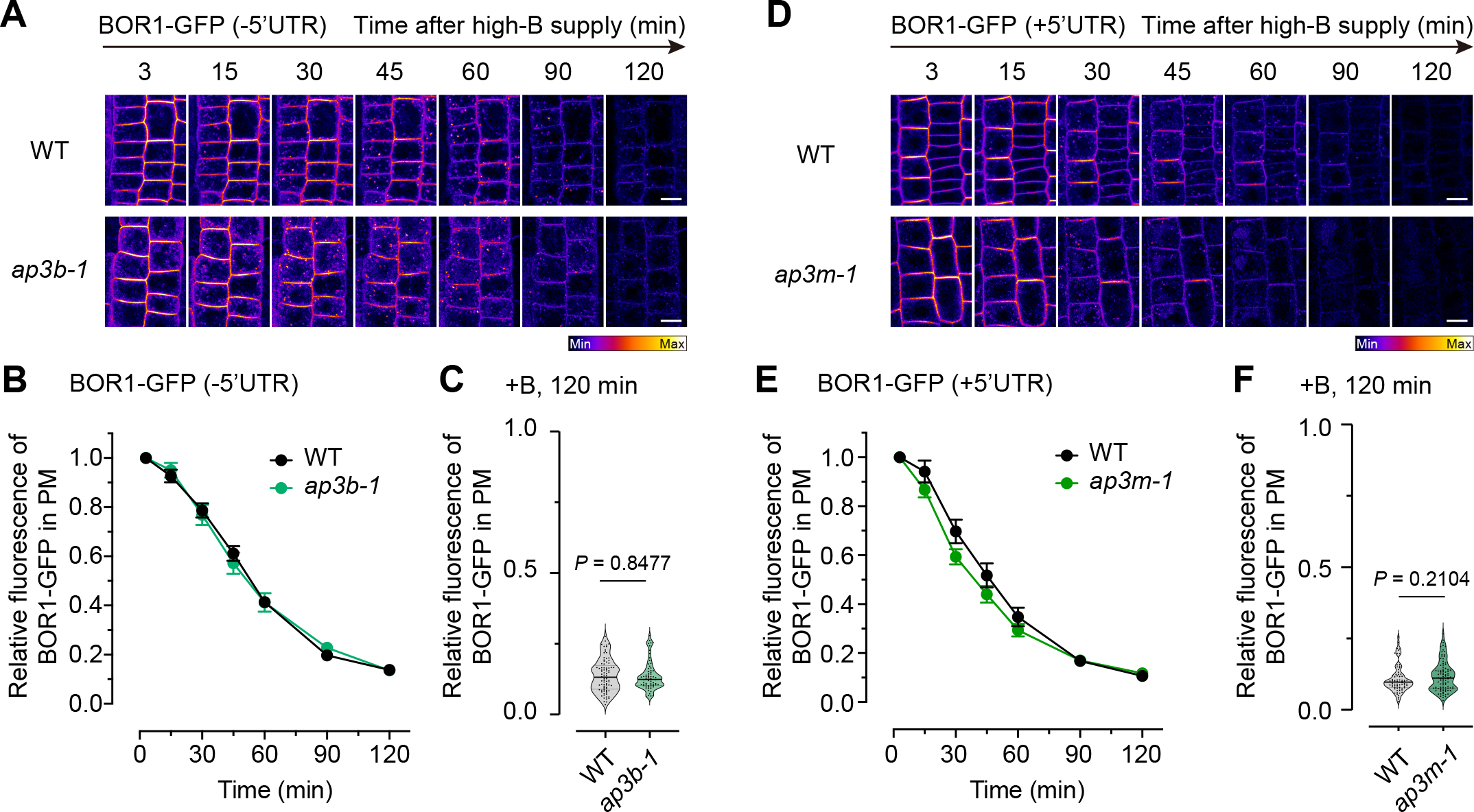
AP-3 is not required for rapid vacuolar sorting of BOR1. Time-lapse imaging of BOR1-GFP in the root epidermis after high-B supply. (A–C) BOR1-GFP with or without 5’UTR which is essential for boron-dependent translational repression in wild-type, *ap3b-1*, and *ap3m-1*. (A, D) Confocal images of BOR1-GFP after high-B (100 μM boric acid) supply. Scale bars represent 5 μm. (B, E) Relative fluorescence of BOR1-GFP in the PM. The relative value was calculated by setting the value at 3 min to 1. *n =* 85 (-5’UTR, WT), 69 (-5’UTR, *ap3b-1*), 74 (+5’UTR, WT), and 94 (+5’UTR, *ap3m-1*) cells from three or four different plants. Data represent mean ± SE. (C and F) Violin plots representing individual data points at 120 min in (B and E). *P* values were determined using the Mann–Whitney *U-*test.

### BOR1 accumulated in the AP-4-positive vacuolar-sorting zone in TGN under high-B conditions

In contrast to AP-3, which is not localized in the TGN, AP-4 has been found within a subdomain of the TGN known as vacuolar-sorting zone (Shimizu et al., 2021; Fuji et al., 2016; Feraru et al., 2010). As the cytoplasmic fraction of BOR1 is also localized in the TGN (Yoshinari et al., 2021b), BOR1 may be localized within the AP-4-positive vacuolar-sorting zone in the TGN. To enable the dynamic analysis of BOR1 and AP4M in living cells, we generated AP4M constructs consisting of genomic fragments, including the promoter and introns, tag red fluorescent protein (TagRFP), and near-infrared fluorescent protein (iRFP). The expression of both AP4M-TagRFP and AP4M-iRFP compensated for the growth of mutants lacking AP4M (Supplementary Fig. S2), indicating that fusion with neither TagRFP nor iRFP compromise the function of AP4M. After shifting to low- and high-B conditions, we performed confocal microscopy of BOR1-GFP and AP4M-TagRFP in root epidermal cells to examine the colocalization of the BOR1 and AP-4 complex (Fig. 4A). BOR1-GFP and AP4M-TagRFP exhibited partial colocalization, irrespective of variations in B concentration (Fig. 4A). Given the challenge of discerning the subdomain of the TGN at the resolution of confocal microscopy, we used super-resolution confocal live-imaging microscopy (SCLIM) to observe transgenic plants expressing BOR1-GFP, AP4M-TagRFP, and the TGN marker iRFP-SYP61 (iRFP-tagged SYNTAXIN OF PLANTS 61). 3D (xyz) imaging using SCLIM revealed that internalized BOR1-GFP colocalized with AP4M-TagRFP after a high-B supply (Figs. 4B and 4C). Through the visualization of colocalized regions of BOR1-GFP and AP4M-TagRFP, we noticed that BOR1 colocalized well with AP4M at approximately 12 min after high-B supply (Figs. 4B, column labeled as “BOR1-GFP in AP4M-TagRFP”). Therefore, the extent of colocalization between BOR1-GFP and AP4M-TagRFP was quantitatively examined by calculating the Pearson’s correlation coefficient (PCC) at 3, 6, 12, and 25 min after high-B supply (Fig. 4D). The PCC between BOR1-GFP and AP4M-TagRFP at 12 min after high-B supply was significantly higher than that at 3 and 6 min, suggesting that BOR1 begins to accumulate in the vacuolar-sorting zone of the TGN within a quarter of an hour after high-B supply. Enlarged images and an intensity profile also evidenced that BOR1-GFP was localized in the AP-4-positive vacuolar-sorting zone of the TGN at 12 min after high-B supply (Figs. 4B and 4C). We also confirmed this notion by observing transgenic plants expressing BOR1-GFP, AP1M2 (AP-1 μ subunit)-TagRFP, and AP4M-iRFP. AP-1 is another TGN-localized AP complex composed of a subdomain that is different from that of AP-4 (Fuji et al., 2016; Shimizu et al., 2021). Consistent with previous studies (Fuji et al., 2016; Shimizu et al., 2021), AP1M2-TagRFP and AP4M-iRFP exhibited mutually exclusive localization (Figs. 4E and 3F). The localization of BOR1-GFP to such an AP4M-labeled subdomain suggests that AP-4 is involved in the sorting of BOR1 (Fig. 4F). Using 3D time-lapse live imaging, we observed that the cytoplasmic fraction of BOR1-GFP gradually increased in volume and colocalized with AP4M-TagRFP (Fig. 5). In line with the findings demonstrating the necessity of AP-4 for the rapid vacuolar transport of BOR1, it can be inferred that AP-4 plays a role in the sequestration of ubiquitinated BOR1 within the vacuolar-sorting zone of the TGN for vacuolar transport.

**Fig. 4.**
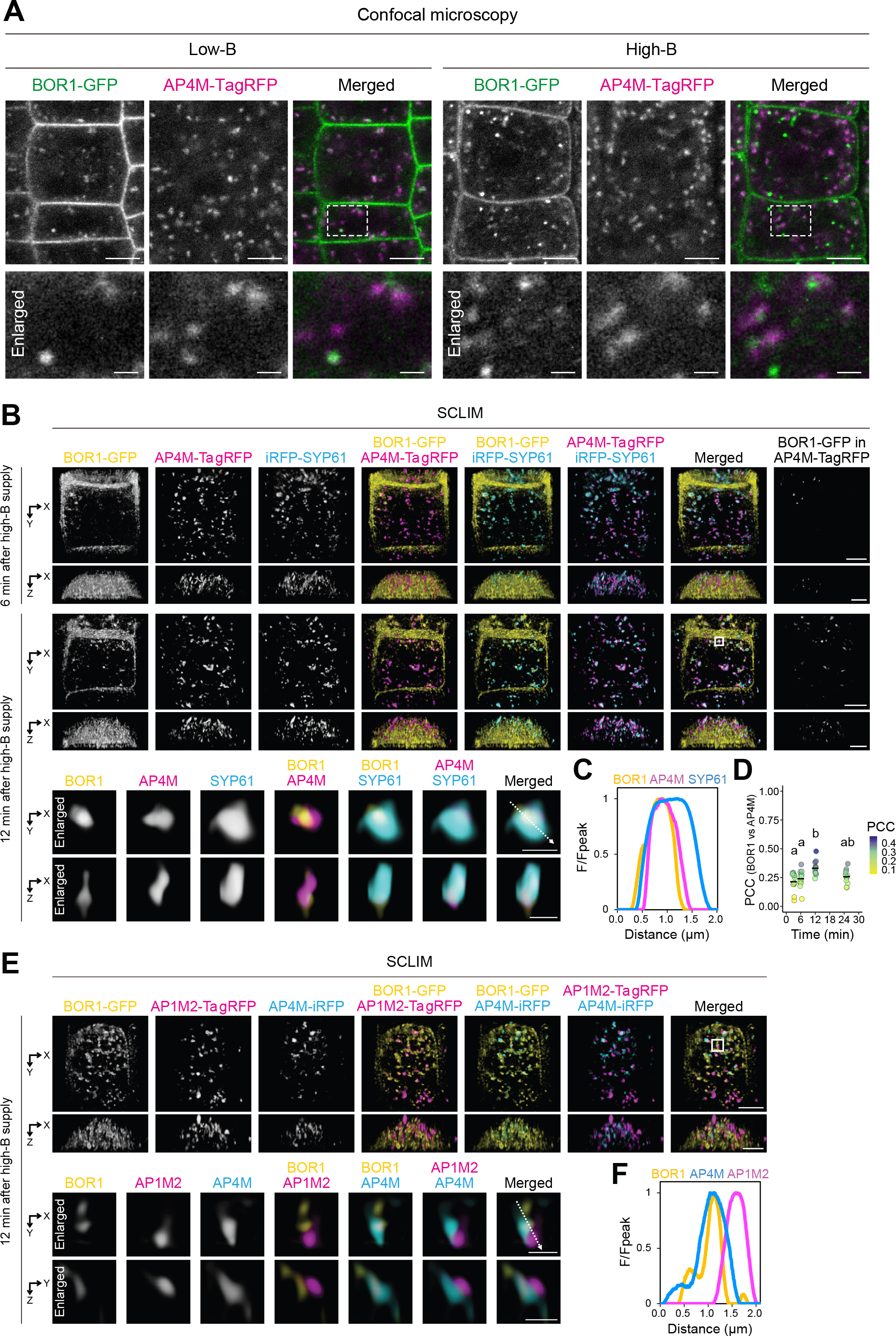
AP-4 partially colocalizes with BOR1 in TGN/EE. (A) Confocal images of BOR1-GFP and AP4M-TagRFP in the root epidermal cells treated with low (0.5 μM) or high (100 μM) concentration of boric acid for 1 h. Enlarged images indicated by boxes with broken lines are shown below. Scale bars represent 5 μm and 1 μm for top and bottom (enlarged image) panels, respectively. (B and E) Top and side views of 3D SCLIM images of BOR1-GFP, AP4M-TagRFP, and iRFP-SYP61 (B) and BOR1-GFP, AP1M2-TagRFP, and AP4M-iRFP (E) in the root epidermal cells after shifting from low (0.5 μM) to high (100 μM) boric acid conditions. Seedlings grown on a medium containing 0.5 μM boric acid were gently put on a coverslip and underlaid with a soft medium containing 100 μM boric acid. Voxels where BOR1-GFP colocalizes with AP4M-TagRFP are visualized in the column labeled as “BOR1-GFP in AP4M-TagRFP.” A representative TGN indicated by the box is magnified and shown below. Scale bars represent 5 μm and 1 μm for lower and higher (enlarged) magnification images, respectively. (C and F) Profiles of relative fluorescence intensity (F/F_peak_) along the arrows in the enlarged image in (B) and (E), respectively. (D) Dot plots representing PCC between BOR1-GFP and AP4M-TagRFP. Bars indicate the mean values. *n =* 12 epidermal cells from four different independent experiments. Different letters denote statistically significant differences from each other (*P* < 0.05; Tukey HSD test in R software).

**Fig. 5.**
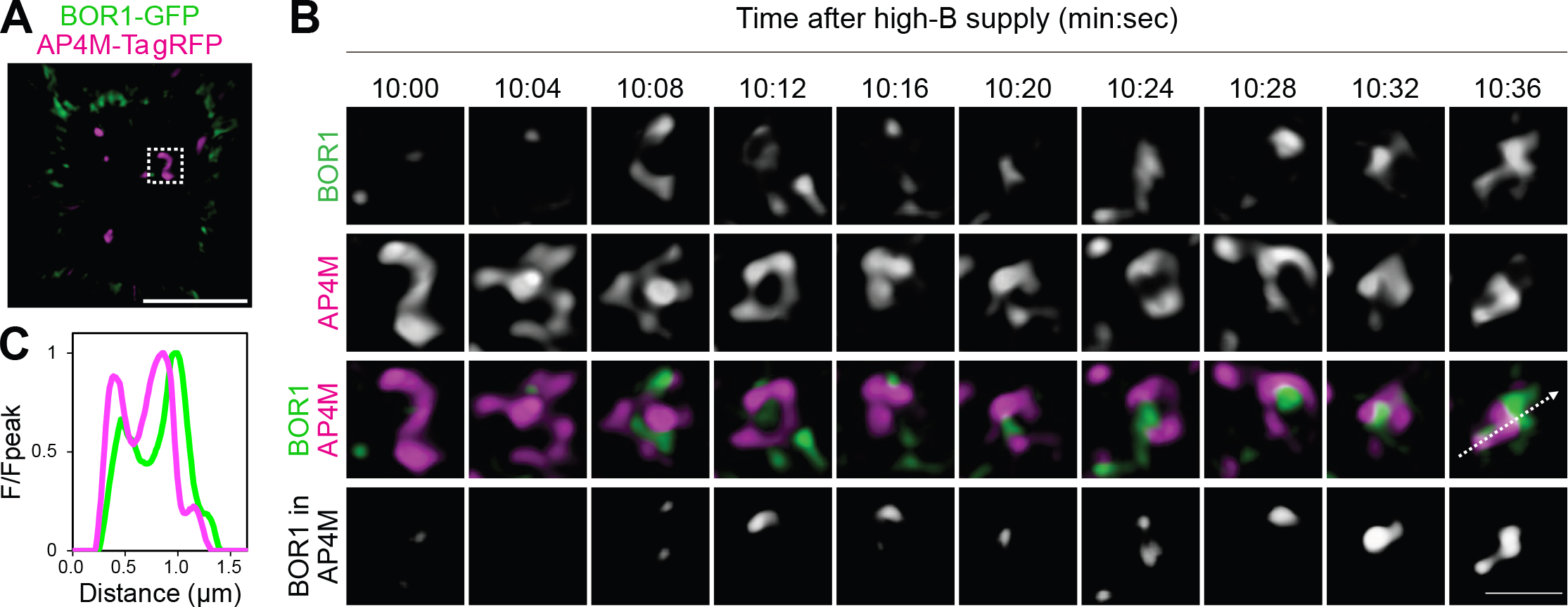
BOR1 gradually accumulates in AP-4-positive compartment. (A) 3D SCLIM images of BOR1-GFP and AP4M-TagRFP in the root epidermal cells 10 min after treatment with high (100 μM) concentration of boric acid. 3D time-lapse images of the AP-4-positive compartment indicated by the box in (A). Scale bars represent 5 μm (A) and 1 μm (B). (C) Profiles of relative fluorescence intensity (F/F_peak_) along the arrows in (B).

## Discussion

To survive in fluctuating environments, plants have evolved a mechanism to rapidly transport the borate transporter BOR1 from the PM to vacuoles upon sufficient B supply in the rhizosphere, with the aim of preventing excessive accumulation of B (Yoshinari and Takano, 2017; Takano and Tanaka, 2023). This process encompasses the polyubiquitination of BOR1 coupled with a B-sensing mechanism during the transport cycle (Yoshinari et al. 2021). Ubiquitinated BOR1 is not recycled back to the PM after endocytosis but is transported to MVBs and vacuoles. In MVBs, BOR1 is located on the membrane of inner vesicles (Viotti et al. 2010). MVBs fuse with the vacuolar membrane and release inner vesicles into the vacuole to degrade the contents, including BOR1. Recent discoveries have elucidated the function of AP-4 in segregating vacuolar cargo within the TGN into the vacuolar-sorting zone (Fuji et al., 2016; Shimizu et al., 2021). In the present study, we found that AP-4 is involved in vacuolar sorting of BOR1. It should be noted that vacuolar sorting of BOR1-GFP was delayed, but still occurred in *ap4m* and *ap4b* mutants (Fig. 2). These results indicate that AP-4 plays a supportive role but is not essential for the B-dependent vacuolar sorting of BOR1. In contrast, a BOR1 mutant with three mutated tyrosine motifs was no longer sorted into MVB after high-B supply (Takano et al., 2010). This conflicting observation implies the necessity for additional factor(s) that recognize tyrosine motifs in the process of endocytosis or MVB sorting. It is likely that AP-4 enhances the rate of MVB sorting of BOR1 by sequestering ubiquitinated BOR1 into the vacuolar sorting zone in the TGN. Previously, we identified that cytoplasmic TOM1-LIKE (TOL) proteins are redundant for the vacuolar sorting of BOR1 (Yoshinari et al., 2018). TOL proteins are ESCRT-0 components required for the initial recognition of ubiquitinated proteins in the Endosomal Sorting Complex Required for Transport (ESCRT) machinery in plant cells (Korbei et al., 2013; Moulinier-Anzola et al., 2020). AP-4 possibly participates in the sequestration mechanism of ubiquitinated BOR1 in the TGN for MVB sorting, alongside TOL proteins.

Our super-resolution imaging of BOR1 and AP4M revealed their precise localization and dynamics in the TGN under high-B conditions (Figs. 4 and 5). We have previously revealed that the TGN harbors at least two distinct subdomains for cargo sorting. One is called the vacuolar-trafficking zone and is composed of AP-4 and R-SNARE VAMP727. The second is the secretory trafficking zone, which is composed of AP-1, clathrin, and VAMP721 (Shimizu et al., 2021). The colocalization of BOR1 and AP4M supports the idea that AP-4 mediates the sorting of BOR1 toward the MVB. This colocalization was observed within 6 min after the high-B supply and became more prominent at 12 min. This finding aligns with previous studies revealing that endocytosed FM4-64 reaches the TGN within 6 min of its uptake (Dettmer et al., 2006) and that BOR1 becomes detectable in the MVB and TGN 30 min after exposure to boric acid (Viotti et al., 2010). Conversely, the findings of these studies suggested that the vacuolar-trafficking zone of the TGN involves an MVB-mediated vacuolar pathway. Although it has been proposed to be generated from the TGN (Scheuring et al., 2011), how cargo proteins are transferred from the TGN to the MVB remains unclear. Time-lapse imaging of BOR1 after high-B supply, in combination with TGN and MVB markers, can be useful for investigating the mechanism of cargo transport between the TGN and MVB in the future. In a previous study, we did not detect physical interactions between the tyrosine motifs of BOR1 and AP1M2 using a Y2H assay (Yoshinari et al., 2019). However, BOR1-GFP was present in the AP4M-labeled vacuolar-trafficking zone and the AP1M2-labeled secretory trafficking zone of the TGN. These observations suggest that AP-1 is involved in the secretory trafficking of BOR1 in a tyrosine motif-independent manner. The secretory trafficking zone of the TGN matures into an intermediate compartment for the secretory trafficking pathway to the PM, known as the Golgi-released independent TGN (GI-TGN)/free TGN (Kang et al., 2011; Uemura et al., 2014; Shimizu et al., 2021). GI-TGN/free TGN consists of, and may produce, a variety of vesicles, including clathrin-coated and non-coated vesicles (Kang et al., 2011; Shimizu et al., 2021). We consider that the presence of BOR1 in the AP1M2-labeled compartments reflects an intermediate state before sorting, and that BOR1 may eventually be sorted into non-coated vesicles for recycling to the PM or relocated to the AP-4-positive vacuolar-trafficking zone.

In this study, we investigated the roles of AP-3 and AP-4 complexes in the trafficking of BOR1. Although we previously demonstrated that the μ subunit of AP-3 complex physically bind the tyrosine motifs of BOR1 by Y2H (Yoshinari et al., 2019), the polar localization and vacuolar sorting of BOR1 was not disturbed in the mutants of AP-3 subunits (Figs. 1A, 1B, and 3). Therefore, the physiological importance of the interactions between BOR1 and AP-3 remains unclear. We consider that the physical interaction ability in the Y2H experiment does not necessarily reflect the actual molecular process *in vivo*. In contrast, we found that AP-4 participates in a conditional vacuolar sorting system during BOR1 trafficking. Specifically, our study highlights the role of AP-4 in the segregation of polyubiquitinated BOR1 within the TGN, facilitating vacuolar sorting of BOR1 under sufficient B conditions. Our findings suggest that plants employ the AP-4-dependent vacuolar sorting system for both the constitutive targeting of functional proteins into the tonoplast or vacuolar lumen (Fuji et al., 2016; Law et al., 2022) and for the rapid degradation of diverse membrane proteins. This rapid degradation process is anticipated to facilitate the optimization of protein abundance in the PM in response to various environmental cues.

## Materials and methods

### Plasmid construction

To construct the AP4M-TagRFP vector (pTH1), *proAP4M:AP4M* genomic sequence in the pENTR/D-TOPO vector (Fuji et al. 2016) was transferred into a binary vector, pGWB659 (Nakamura et al., 2010), using the LR reaction of the Gateway system (Invitrogen). To construct the AP4M-iRFP vector (pTH4), the pTH1 vector was digested with BamHI and SacI, and the iRFP coding sequence, amplified using the primers, described in Supplementary Table S1, was inserted using the in-fusion cloning technique (Clontech).

### Plant materials

*Arabidopsis thaliana* Col-0 was used as the wild-type. We used KK8-5 line (*bor1-1* background) for *BOR1pro:BOR1-GFP (+5’UTR)* (Yoshinari et al., 2019, 2021a). For *BOR1pro:BOR1-GFP (-5’UTR)* plants, binary vector pTH3 (Aibara et al., 2018) was introduced into *bor1-1*. Transgenic plants expressing *SYP61pro:iRFP713-SYP61* were previously generated (Shimizu et al., 2021). Seeds of the T-DNA insertion lines *ap3b-1* (*pat2-2*, SAIL_1258_G03, Niihama et al., 2009; Feraru et al., 2010) and *ap4b-2* (*gfs4-3*, SAIL_796_A10, Fuji et al., 2016) were provided by the Arabidopsis Bioresource Center (ABRC). Seeds of *ap3m-1* (SALK_015332; Niihama et al. 2009) were provided by Dr. Miyo Terao Morita (National Institute for Basic Biology, Japan). Seeds of *ap4m-3* (*gfs5-3*, SALK_014326, Fuji et al., 2016) and *ap4m-4* (*gfs5-4*, SALK_044748, Fuji et al., 2016) were provided by Dr. Tomoo Shimada (Kyoto University). *BOR1-GFP/ap3m-1, BOR1-GFP/ap4m-3, BOR1-GFP/4m-4*, and *BOR1-GFP/ap4b-1* were generated by crossing. *BOR1-GFP/ap3b-1* was generated by the direct transformation of *ap3b-1* plants with pTH3. Genotyping PCR was performed using the primers listed in Supplementary Table S1. To generate plants co-expressing BOR1-GFP and AP1M2, AP4M, or SYP61 fused with fluorescent proteins, *AP1M2pro:AP1M2genomic-TagRFP, AP4Mpro:AP4Mgenomic-TagRFP, AP4Mpro:AP4M-iRFP*, and/or *SYP61pro:iRFP713-SYP61* were introduced into KK8-5 via Agrobacterium-mediated floral-dip transformation (Clough and Bent, 1998). For complementation analysis using AP4M-iRFP and AP4M-TagRFP, the *AP4Mpro:AP4Mgenomic-TagRFP* and *AP4Mpro:AP4M-iRFP* constructs were introduced into the knockout alleles of *ap4m, gfs5-1* and *gfs5-2* (Fuji et al., 2016).

### Plant growth condition

For imaging analysis, plants were grown on low-B MGRL medium containing 0.5 μM boric acid, 1.5% (w/v) gellan gum (Wako Pure Chemicals, Osaka, Japan), and 1% (w/v) sucrose for 4 or 5 d. Seeds were surface-sterilized by 70% ethanol and washed 4 times with sterilized ultra-pure water (Milli-Q water, Merck-Millipore) followed by stratification at 4 ºC for 2 d. Subsequently, the seeds were sown onto the surface of low-B MGRL medium and incubated vertically in the plant growth chamber BioTRON (NK system, Osaka, Japan) under 16-h/8-h light/dark cycles. For complementation analysis, plants were grown on MGRL medium containing 30 μM boric acid, 1%(w/v) gellan gum, and 1%(w/v) sucrose for 7 d.

### Confocal microscopy

For confocal laser scanning microscopy, we used a Zeiss LSM800 (Carl Zeiss, Oberkochen, Germany) equipped with 40x water-immersion (1.1 NA LD C-Apochromat, Zeiss), 63x oil-immersion lenses (1.4 NA Plan-Apochromat, Zeiss) and 488 and 561-nm lasers, and a Leica TCS-SP8 gSTED (Leica Microsystems, Wetzlar, Germany) equipped with 100x oil-immersion objective (HL PLAN APO CS2 1.40 oil-immersion, Leica Microsystems) and a white-light laser (used as a light source). Room temperature was adjusted at 23 ºC during all the steps including sample preparation and imaging. To measure BOR1-GFP fluorescence after a high-B supply, a Zeiss LSM800 confocal microscope was used with a detection window of 500–530 nm. For colocalization analysis of BOR1-GFP and AP4M-TagRFP, a Leica TCS-SP8 gSTED confocal microscope was used with excitation/detection wavelengths of 488/495–540 and 561/570–650 nm for GFP and TagRFP, respectively.

### Calculation of BOR1-GFP polarity index

The polarity index of BOR1-GFP was calculated as previously described (Yoshinari and Takano, 2020). In brief, the fluorescence intensity of BOR1-GFP was measured along the radial PM, and the total intensity of the inner half was divided by that of the outer half of each cell. For efficient quantification, we used a modified Python code based on the polarityindexcalculator (https://github.com/totti0223/polarity_index_calculator) (Yoshinari et al., 2021b). The modified code is described in the Supplementary Methods.

### Timelapse imaging of BOR1-GFP degradation

Time-lapsed imaging was performed as previously described (Yoshinari and Takano, 2020). Briefly, 4-day-old seedlings were transferred on a droplet of MGRL liquid medium containing 100 μM B on glass-bottom dish and covered by MGRL soft medium containing 100 μM B. Fluorescence of BOR1-GFP in the root epidermal cells was acquired by confocal microscopy. The focus was adjusted manually during imaging.

### Super-resolution confocal live imaging microscopy (SCLIM)

For super-resolution imaging, the SCLIM developed at RIKEN was used as previously described, with minor modifications (Kurokawa and Nakano, 2020; Shimizu et al., 2021). The SCLIM system consists of: an Olympus IX73 inverted microscope (Olympus, Tokyo, Japan) equipped with a 100x silicone-immersion objective (NA 1.35 UPLSAPO100XS); a custom-made spinning-disk confocal scanner (Yokogawa Electric, Tokyo, Japan); a custom-made spectroscopic unit (detection wavelengths = 490–545 nm for GFP; 580–660 nm for TagRFP; 680 nm– for iRFP); a custom-made image intensifier system (Hamamatsu Photonics, Hamamatsu, Japan); 4x and 2/3x intermediate lenses (Nikon, Tokyo, Japan); three EM-CCD cameras (Hamamatsu Photonics, Hamamatsu, Japan); and three solid-state excitation lasers with emission at 473 nm (Cobolt, Stockholm, Sweden), 561 nm (Cobolt, Stockholm, Sweden), and 671 nm (CrystaLaser, Nevada, USA). The samples were prepared as described above. For 3D (xyz) images, we collected 71 optical sections spaced 0.1 μm apart (z-range = 7 μm). For 3D time-lapse images, we collected 18 optical sections spaced 0.3 μm apart (z-range = 5.4 μm). Z-stack images were reconstructed into 3D images and processed using deconvolution algorithms with theoretical point-spread functions in Volocity software (Quorum Technologies, East Sussex, UK). Regions of the epidermal cells were cropped manually and displayed to remove the fluorescence signal from the lateral root cap cells. Visualization of colocalized regions of BOR1-GFP and AP4M-TagRFP was performed using the Volocity software function “Generate Colocalization, PDM channels,” which generate the colocalized region if voxel values for both channels are above the mean of each channel. The PCC was calculated from the area of the epidermal cell using Volocity.

## Acknowledgments

We thank Dr. Tomoo Shimada (Kyoto University) for providing AP4M_TOPO vector and seeds of, *gfs5-1, gfs5-2, gfs5-3* (*ap4m-3*), *gfs5-4* (*ap4m-4*), and *ap4b-2* (*gfs4-3*), Dr. Miyo Terao Morita (National Institute for Basic Biology) for seeds of *ap3m-1*, Dr. Kyoko Miwa (Hokkaido University) for pTH3 vector, and Dr. Tsuyoshi Nakagawa (Shimane University) for pGBW659 vector. We also thank Kazue Kanokogi (Osaka Metropolitan University), Yuka Ohkita (Osaka Metropolitan University), Kayo Konishi (Hokkaido University), Tomoko Shimizu (Hokkaido University), Yuko Kuroda (Nagoya University), and Yuki Ichikawa (Nagoya University) for their excellent technical assistances.

## Author contributions

J.T., A.Y. and T.U. conceptualized the project. A.Y., Y.S., T.U., and J.T. designed the experiments. A.Y., Y.S., and T.H. performed the experiments. A.Y. and Y.S. performed visualization. A.Y., Y.S., T.H., and J.T. acquired essential materials.

A.Y. and J.T. wrote the original draft. A.Y., Y.S., A.N., T.U., and J.T. reviewed and edited the manuscript. A.Y., T.U., and J.T. acquired funding, and J.T. administered the project.

