## Supplementary material for "Rapid Vacuolar Sorting of the Borate Transporter BOR1 Requires the Adaptor Protein Complex AP-4 in Arabidopsis": Supplmentary Figures

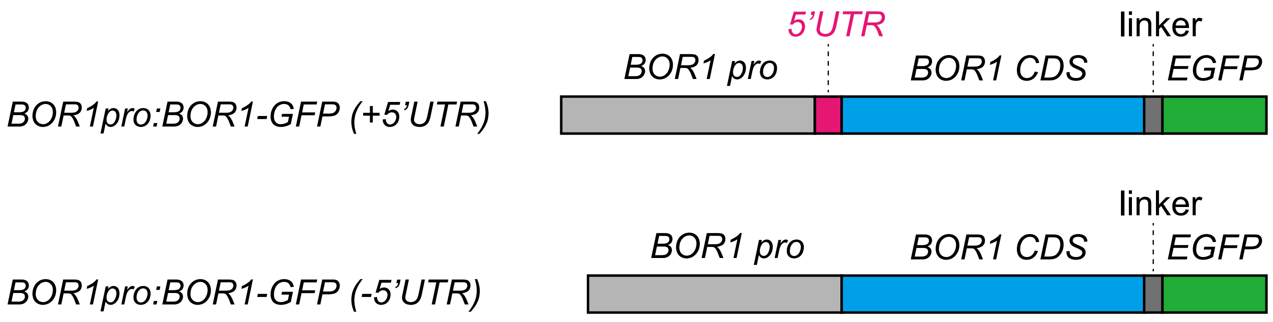


**Supplementary Fig. S1. Structure of *BOR1-GFP* transgene**

Schematic illustration of *BOR1pro:BOR1-GFP (±5' UTR)*. BOR1 pro, the own promoter of BOR1; BOR CDS, protein-coding sequence of BOR1; EGFP, enhanced green fluorescent protein; Linker, linker sequence consisting of Gly-Gly-Gly-Gly-Ala (5 amino acid residues).

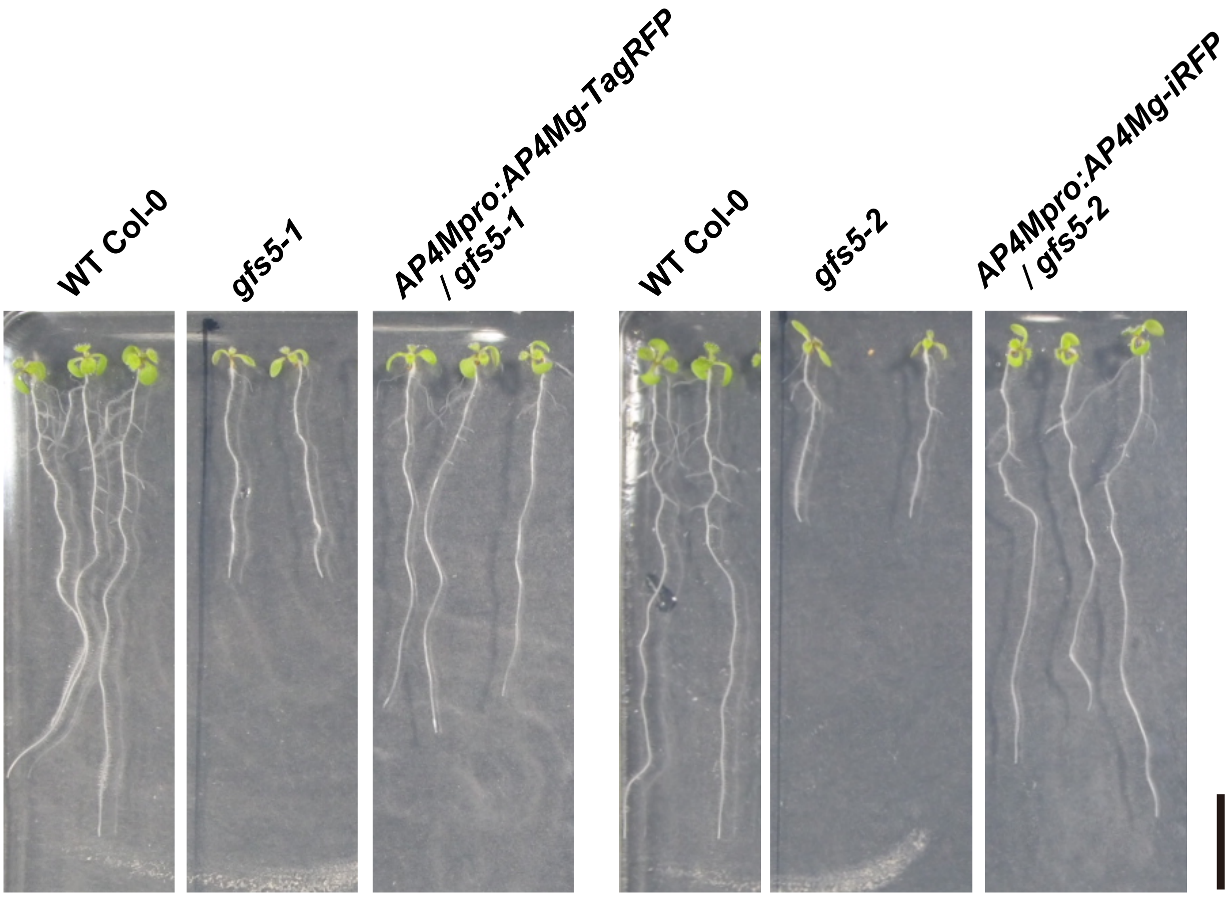


**Supplementary Fig. S2. Genetic complementation of growth of *ap4m* mutants by AP4M-TagRFP and AP4M-iRFP**

*AP4Mpro:AP4Mgenomic-TagRFP/gfs5-1* and *AP4Mpro:AP4M-iRFP/gfs5-2* were grown in modified MGRL medium containing 30 μM B for 7 d. The scale bars indicate 10 mm.
