## Supplementary Table 1 for "Rapid Vacuolar Sorting of the Borate Transporter BOR1 Requires the Adaptor Protein Complex AP-4 in Arabidopsis"

### Supplementary Table S1. Primers used in the study

| Name | Sequence (5' to 3') | Purpose |
|---|---|---|
| SAIL_1258_G03 LP | CATCGGAAGTGAAGAAGCTTG | Genotyping of ap3b-1 (pat2-2, SAIL_1258_G03) |
| SAIL_1258_G03 RP | CCAGCTGCTGCTAGTACAACC | |
| ap3*μ*-1 LP | TTGTCTGAGTACCTTGGTGGG | Genotyping of ap3m-1 (SALK_015332) |
| ap3*μ*-1 RP | CTGGAAAACCAGTGAGCTGAG | |
| SALK_014326_LP | CGTTTTATTGCTACAGACCGG | Genotyping of *ap4m-3* (*gfs5-3*) |
| SALK_014326_RP | CTTCATTTGAATGGTGCCATC | |
| SALK_044748 LP | TGAGACTCCCTCACCAACATC | Genotyping of *ap4m-4* (*gfs5-4)* |
| SALK_044748 RP | TCAAGGAATCCAACAAAATGC | |
| LP ap4β SAIL_796_A10 | CATTGTGGTGTTTCACTGCAG | Genotyping of ap4b-2 (gfs4-343, SAIL_796_A10) |
| RP ap4β SAIL_796_A10 | TGGCTCCATTGTGGATACTTC | |
| LBb1.3 | ATTTTGCCGATTTCGGAAC | Genotyping for SALK lines |
| LB for SAIL | TAGCATCTGAATTTCATAACCAATCTCGATACAC | Genotyping for SAIL lines |
| iRFP_fw | gttgataacagcATGGCTGAAGGATCCGTCGCC | Cloning of AP4M-iRFP |
| iRFP_rv | gatcggggaaattcgTCACTCTTCCATCACGCCGATCTG | |
