## Supplementary Method for "Rapid Vacuolar Sorting of the Borate Transporter BOR1 Requires the Adaptor Protein Complex AP-4 in Arabidopsis"

**Modified Python code of Polarity Index Calculator**

from ij import IJ

from ij.plugin.frame import RoiManager

from ij.plugin.filter import ParticleAnalyzer, BackgroundSubtracter, EDM

from ij.measure import ResultsTable

from ij.gui import ProfilePlot

from ij.io import DirectoryChooser

import csv

import os

\# specify path to csv file you want to generate

\# ex. f = open(PATH, "wb")

\# bellow is an example. replace the path to your needs.

\# Get the image filename

imp = IJ.getImage()

img_filename = imp.getTitle()

\# Create the output file path using the image filename

PATH = os.path.join(os.path.dirname(imp.getOriginalFileInfo().directory),

"{}_result.csv".format(os.path.splitext(img_filename)[0]))

###############

assert PATH.endswith(".csv"), "you must specify a csv in PATH"

RM = RoiManager()

rm = RM.getRoiManager()

numROI = rm.count

assert numROI != 0, "you need a ROI added to ROI manager to run this script"

f = open(PATH,"wb")

writer = csv.writer(f)

imp = IJ.getImage()

writer.writerow(["ROI_No","ROI_Length","PMin","PMout","PI"])

profiles = []

for i in range(numROI):

rm.select(i)

pf = ProfilePlot(imp)

profile = pf.getProfile()

length = len(profile)

PMin = sum(profile[:length//2])

PMout = sum(profile[length//2:])

PI = PMin / PMout

writer.writerow([str(i+1).zfill(3),len(profile),PMin,PMout,PI])

#profiles.append(list(range(len(profile))))

row = ["ROI_No:"+str(i+1)]

row.extend(profile)

profiles.append(row)

writer.writerow([""])

writer.writerow(["Raw Data"])

for i,row in enumerate(profiles):

writer.writerow(row)

f.close()

print("Done. Result saved in the bellow path")

print(PATH)

###############
